# ParaSurf: A Surface-Based Deep Learning Approach for Paratope-Antigen Interaction Prediction

**DOI:** 10.1101/2024.12.16.628621

**Authors:** Angelos-Michael Papadopoulos, Apostolos Axenopoulos, Anastasia Iatrou, Kostas Stamatopoulos, Federico Alvarez, Petros Daras

## Abstract

**Motivation:** Identifying antibody binding sites, is crucial for developing vaccines and therapeutic antibodies, processes that are time-consuming and costly. Accurate prediction of the paratope’s binding site can speed up the development by improving our understanding of antibody-antigen interactions.

**Results:** We present ParaSurf, a deep learning model that significantly enhances paratope prediction by incorporating both surface geometric and non-geometric factors. Trained and tested on three prominent antibody-antigen benchmarks, ParaSurf achieves state-of-the-art results across nearly all metrics. Unlike models restricted to the variable region, ParaSurf demonstrates the ability to accurately predict binding scores across the entire Fab region of the antibody. Additionally, we conducted an extensive analysis using the largest of the three datasets employed, focusing on three key components: (1) a detailed evaluation of paratope prediction for each Complementarity-Determining Region loop, (2) the performance of models trained exclusively on the heavy chain, and (3) the results of training models solely on the light chain without incorporating data from the heavy chain.

**Availability and Implementation:** Source code for ParaSurf, along with the datasets used, preprocessing pipeline, and trained model weights, are freely available at https://github.com/aggelos-michael-papadopoulos/ParaSurf.

**Supplementary information:** Supplementary data are available at *Bioinformatics* online.

## Introduction

Antibodies, also known as immunoglobulins, are crucial components of the immune system that specifically recognize and neutralize foreign molecules (antigens) such as pathogens and toxins. Structurally, antibodies are Y-shaped proteins, composed of two identical heavy chains and two identical light chains. The variable regions (V) of both chains form the antigen-binding fragment (Fab domain), while the constant region (Fc domain) plays a pivotal role in immune effector functions. Within the Fab region, the variable domain (Fv) houses the complementarity-determining regions (CDRs), which are hypervariable loops responsible for the high specificity of antigen binding and the framework residues. These CDR loops, particularly CDR3, form the key interface for antigen binding (Janeway Jr et al. 2001). The ability of antibodies to precisely bind antigens ensures a targeted immune response, facilitating antigen neutralization and the recruitment of other immune cells.

Studying antibody-antigen (Ab-Ag) interactions is critical for understanding immune recognition and developing therapeutic targets. Structural biology techniques such as X ray crystallography (Smyth and Martin 2000) and Nuclear Magnetic Resonance (NMR) (Rhodes 2017) have historically been used to determine high-resolution structures of antibody-antigen complexes. X-ray crystallography provides detailed atomic-resolution structures, while NMR can capture more dynamic aspects of the interaction in solution. Modern methods, including cryo-electron microscopy (cryo-EM) (Vant et al. 2022) and biophysical techniques such as surface plasmon resonance (SPR) (Vant et al. 2022), complement these approaches by providing real-time interaction data and structural information without the need for crystallization. Together, these techniques offer comprehensive insights into how antibodies recognize and neutralize antigens, guiding the design of vaccines and antibody-based therapies.

These traditional methods, while effective, are time-consuming, expensive and often not scalable to the increasing demand of high-throughput data in immunology research. This has led to a shift toward computational approaches, particularly in the field of deep learning. Various models have emerged to address this problem with different strategies. For example (Cohen et al. 2023)is a method that integrates deep learning with X-ray scattering data is presented to resolve the structure of antibody-antigen complexes)

The revolutionary achievements in protein structure prediction - 3D folding, led by Alphafold (Jumper et al. 2021) have shifted the focus of the scientific community toward predicting the 3D structure of antibodies. With the success of Alphafold, several new methods have appeared specifically for antibody structure prediction, such as ABodyBuilder3 (Kenlay et al. 2024), offering reliably predicted structures of antibodies. The recent release of Alphafold3 (Abramson et al. 2024) has further enhanced the accessibility of predicting complex structures, including antibodies. In addition, the development of Antifold (Høie et al. 2023), an inverse folding approach, enables the efficient design of antibody sequences that preserve structural integrity. This ensures that crucial binding regions like the CDR loops are optimized without disrupting the overall protein fold. Antifold accelerates antibody development by predicting mutations that improve binding affinity and stability, reducing the need for experimental trial and error. Advances like AlphaFold provide accurate 3D structure predictions, offering deeper insights into antibody domains and configurations.

While breakthroughs like AlphaFold have revolutionized the prediction of 3D structures, general protein-protein interaction (PPI) methods, such as MaSIF (Gainza et al. 2020), BipSpi (Sanchez-Garcia et al. 2019), DIPS (Townshend et al. 2019), ProteinMAE (Yuan et al. 2023), and DockNet (Williams et al. 2023), have managed to tackle the binding interface prediction task in various PPI contexts. These methods focus on identifying interaction sites across a wide range of protein complexes, offering general solutions to PPI tasks.

However, to gain a thorough understanding of the paratope-antigen (or receptor-antigen) interactions, it is crucial to develop specialized approaches that focus on predicting the binding site of the paratope. This is essential for accurately mapping antibody-antigen interactions, thus improving our understanding of immune responses and guiding further research in immunology. For instance, Parapred (Liberis et al. 2018) achieved notable results in predicting paratope binding sites by focusing solely on the antibody sequence. It combined Convolutional Neural Networks (CNNs) to capture local features with Long Short-Term Memory (LSTM) cells to model long-range dependencies across the entire sequence, outperforming earlier methods (Krawczyk et al. 2013, Tsuchiya and Mizuguchi 2016).

Another approach, introduced by Daberdaku (Daberdaku and Ferrari 2019), utilized 3D Zernike Descriptors to predict the antibody-antigen binding interfaces. By capturing both geometric and physico-chemical properties of the paratope surface, this method managed to classify surface patches as binding or non-binding using a Support Vector Machine (SVM). The combination of rotationally invariant descriptors and SVM allowed for accurate prediction of binding interfaces.

PECAN (Pittala and Bailey-Kellogg 2020) introduced a deep learning framework designed to predict paratope-antigen binding interfaces using Graph Neural Networks (GNNs). PECAN captured the spatial relationships between residues and incorporated an attention layer to account for the binding context provided by the partner protein. Combining these techniques with transfer learning from the general protein-protein interaction data Docking Benchmark Version 5, DBv5 (Vreven et al. 2015), PECAN emerged as a competitive deep learning approach for paratope prediction. Additionally, Daberdaku released an antibody-antigen dataset, which, with some exclusions by PECAN (11 complexes), is widely regarded as the prominent benchmark dataset for paratope prediction, consisting of 460 antibody-antigen complexes.

On the same note, Paragraph (Chinery et al. 2023), the current state-of-art method at this paratope prediction benchmark, also employed GNNs. Unlike previous models, Paragraph required only the modeled structure of the antibody and did not depend on antigen information, making it highly versatile. The model applied equivariant GNN layers to capture spatial relationships between residues and used simple feature vectors for each residue. To further boost Paragraph’s performance the authors created a new expanded dataset consisting of 1,086 paratope-antigen complexes and validated the model with improved evaluation metrics.

Lastly, MIPE (Wang et al. 2024) introduced a novel approach to paratope and epitope prediction by using multi-modal contrastive learning and interaction informativeness estimation. MIPE integrated both sequence and structure data of antibodies and antigens, leveraging multi-modal contrastive learning to maximize the representations of binding and non-binding residues across different modalities. MIPE also validated its method on a new dataset consisting of 626 antibody-antigen complexes, achieving state-of-the-art results, when compared to previous methods PesTo (Krapp et al. 2023) and AG-Fast-Parapred (Deac et al. 2019) and Paragraph on this specific dataset.

In this work, ParaSurf is introduced as a novel deep learning approach inspired by DeepSurf (Mylonas et al. 2021). DeepSurf leveraged surface-derived features from the protein’s solvent-accessible area to predict the binding sites on the protein where it interacts with its corresponding ligand (small molecule). Building upon this concept, ParaSurf makes extracts geometric, chemical and electrostatic force-field features from the paratope’s surface representation and then train a hybrid architecture consisting of a 3D ResNet and a transformer layer. This design allows ParaSurf to achieve state-of-the-art performance across nearly all metrics on three key antibody-antigen benchmarks: the well-established PECAN dataset (460 complexes), the Paragraph expanded dataset (1,086 complexes), and the MIPE dataset (626 complexes).

Similar to the Paragraph model, ParaSurf is also an antigen-agnostic model, meaning it does not require antigen information during training. However, unlike Paragraph, ParaSurf extends its predictive power beyond the Fv region, covering the entire Fab domain of the antibody, thus addressing the inherent class imbalance nature of the paratope prediction task. Additionally, ParaSurf performs extensive analysis on each of the CDR loops, on the CDR plus two extra residues on either end (CDR±2) regions, on the Fv region and the Fab region. Additionally, using the expanded dataset, ParaSurf undergoes specialized retraining experiments on individual antibody chains, focusing separately on the heavy chain and the light chain. The CDR loops were considered using the IMGT numbering rule (Lefranc et al. 2009) as in Paragraph.

In summary, the main contributions of this paper are described as follows:

- **Field-force incorporation**: ParaSurf incorporates novel features, including geometrical, chemical, and electrostatic information, within a hybrid 3D ResNet and transformer architecture.
- **Paratope generalization:** ParaSurf provides a generalized approach, predicting with remarkable accuracy the binding scores of each residue across the entire Fab region, rather than being restricted solely to the variable (Fv) or to the CDR±2 region.
- **Extensive analysis:** Concerning the Paragraph-expanded dataset, ParaSurf delivers a detailed analysis on each of the CDR loops and the framework region. It also includes additional retraining experiments focusing on each individual antibody chain of the paratope (heavy chain and light chain), showcasing state-of-the-art performance.
- **State-of-the-art performance:** ParaSurf achieves superior results across three major antibody-antigen benchmarks; PECAN (460 complexes), Paragraph expanded dataset (1,086 complexes), and MIPE dataset (626 complexes). It consistently delivers state-of-the-art performance on the most critical metrics, AUC-ROC and AUC-PR, across all three datasets. Additionally, ParaSurf excels in predicting binding sites on the most variable CDR3 regions, for both the heavy and light chains, and achieves leading performance on the framework region.
- **Reproducibility:** Open-source code, datasets, and trained model weights for each training scenario are available for public use, encouraging further research and development in the field at https://github.com/aggelos-michael-papadopoulos/ParaSurf.

### Data

To allow direct comparison with the previously published methods ParaSurf was trained and tested on exactly the same datasets.

#### PECAN dataset

This dataset is widely used as the benchmark for comparing the performance of most paratope prediction methods. ParaSurf was trained and tested on the same complexes used by PECAN. It consists of 460 paratope-antigen complexes, with 205 designated for training, 103 for validation, and 152 for testing. This separation was made using CD-HIT (Li and Godzik 2006)to ensure that paratopes shared no more than 95% pairwise sequence identity. It includes only complexes with paired heavy and light chains, with a resolution below 3°A and protein antigens. Subsequent datasets, such as the Paragraph Expanded and MIPE datasets, followed the same data separation technique established by the PECAN dataset.

#### Paragraph expanded dataset

The authors from Paragraph also trained their model on a new expanded dataset derived from the Structural Antibody Database, SAbDab (Schneider et al. 2022) on March 31, 2022. This dataset contains 1,086 antibody-antigen complexes. To ensure consistency with the original work, we used the exact same data split as Paragraph, dividing the dataset into 60% for training, 20% for validation, and 20% for testing, without making any additional modifications or random splits.

#### MIPE dataset

This dataset is also derived from SAbDab. It consists of 626 antibody-antigen pairs. Following the same principles as the original MIPE method, we randomly split the dataset, allocating 90% for cross-validation (CV) and 10% as an independent test set. We implemented a 5-fold CV, where the model was trained on each fold and tested on the same independent test set, as MIPE authors suggest.

ParaSurf was trained and tested using five different distinct training schemes to ensure extensive performance analysis. These include 1) training on the PECAN dataset, 2) training on the Paragraph expanded dataset, 3) training on the expanded dataset using only the heavy chain, 4) training on the expanded dataset using only the light chain, and 5) training on the MIPE dataset. The weights for each training scheme, along with the specific splits for each dataset are available at https://github.com/aggelos-michael-papadopoulos/ParaSurf.

## Proposed method

In line with previous methods, a residue is considered part of the paratope’s binding site if any of its heavy atoms (non-hydrogen atoms) is located within **4.5°A** of any antigen-heavy atom.

### Data preprocessing

The pre-processing phase of ParaSurf begins by refining the antibody-antigen complex. Specifically, we remove non-essential components such as water molecules, ions, and ligands from the receptor’s PDB structure. Then, we perform a sanity check to ensure that each antibody-antigen pair meets the 4.5°A Euclidean distance criterion. If a pair fails to meet this condition, it is excluded from the dataset. This step is particularly important in training scenarios 3 and 4, where the detachment of one chain may cause some receptor-antigen pairs to violate this criterion, resulting in their removal (Supplementary Information).

After the initial filtering, we use the DMS software (UCSF 2024) to generate the solvent-accessible surface (SAS) representation of the paratope, which resembles the van der Waals surface of the molecule. Along with the receptor’s surface, we obtain the corresponding normal vectors for each surface point, pointing outward from the surface. To ensure sufficient surface point coverage, we set the density value to 0.5, which is recommended for large molecules like antibodies (UCSF 2024). Next, to deal with the class imbalance nature of the antibody-antigen task we perform a balanced sampling. For each receptor, we select 800 positive surface points (those ¡4.5°A from the antigen) and 800 negative surface points, randomly selected from the surface of the receptor (those ¿4.5°A away from the antigen). Suppose fewer than 800 positive atoms are present on the paratope’s binding surface. In that case, we select as many as are available and adjust the number of negative atoms to match, ensuring a balanced sample. For example, if only 500 binding surface points can be gathered, we will randomly select 500 non-binding surface points from the receptor’s surface.

#### Feature extraction

Once we have selected a set of 1,600 surface points, we proceed to the feature extraction phase. For each surface point, a local grid of size 41×41×41 with a voxel resolution of 1°A is created, following the recommendation from DIPS (Townshend et al. 2019) for protein-protein interactions. The grid is oriented perpendicular to the surface, similar to DeepSurf’s approach, which aligns the grid based on the normal vectors of the surface points. This alignment helps to alleviate the rotation-sensitivity problem inherent in voxelized representations. Each voxel grid is populated with 22 features calculated per atom, describing the atom and its surrounding environment. These features are:

- **18 chemical features** introduced by (Stepniewska-Dziubinska et al. 2018), including 9 atom classes (B, C, N, O, P, S, Se, halogen and metal), 4 atom properties (hybridization, heavy valence, heterovalence, partial charge), and 5 SMARTS-based (a language for describing molecular patterns) features such as hydrophobicity, aromaticity, acceptor/donor properties, and ring structures, same as DeepSurf (Mylonas et al. 2021).
- **2 electrostatic force field features along with 2 atom radii** derived from pdb2pqr (Dolinsky et al. 2004). These include AMBER (Assisted Model Building with Energy Refinement) (Weiner and Kollman 1981), a force field used to simulate the interactions between atoms in molecular systems and CHARMM, another widely used force field that provides a detailed mathematical model of the forces between atoms in biological molecules.

Overall we obtain a combination of geometric, chemical and Electrostatic Force-Field features. Geometric features, capture the shape and orientation of atoms on the molecular surface, using the van der Waals surface and grid-based sampling. Chemical features provide atomic-level properties and functional group characteristics, which are crucial for recognizing the types of interactions possible between atoms. Electrostatic Force-Field features quantify interaction energies through electrostatic potentials, adding an energy-based perspective to the surface information from DMS. These combined features aid the prediction of the binding affinities and the stability of antibody-antigen interactions.

Each surface point contributes a 4D tensor of size 41×41×41×22, which is then used as an input to ParaSurf’s deep neural network. The 41×41×41 grid, with a voxel resolution of 1°A, ensures that all atoms within a 20°A radius from each reference surface point are included, capturing the local molecular geometry alongside chemical and electrostatic properties. Before the training round, these tensors are randomly rotated by 90 degrees along one of the three axes to further reduce rotation sensitivity.

The antibody-antigen binding site prediction task is treated as a binary classification problem, where each residue—comprising all its atoms—is labeled as either binding or non-binding. If an atom’s predicted score exceeds 0.5 threshold, it is classified as part of the binding site. After predicting the binding score for each surface point, we assign a residue-level score by taking the maximum score out of all surface point that belong to a particular residue, according to Equation 1. This threshold is consistent with previous methods, allowing for direct comparisons.

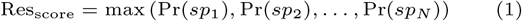

where Pr(*sp*_*i*_) is the prediction score for the surface point *i*. If Res_score_ *>* 0.5, then the residue belongs to the binding site.

Residue-level predictions are then compared to ground truth labels, where binding residues are defined based on the 4.5°A distance threshold. This validation method ensures accurate residue-level predictions and allows for detailed comparison with the ground truth labels. The above feature extraction pipeline is summarized in algorithm 1

In the top section of Figure 1 (Feature Extraction), we depict the feature extraction process for an antibody-antigen complex (PDB code “5EOC”) from the PECAN test set. Both Fab and Fv regions of the paratope are visualized. This section illustrates the creation of the 41×41×41×22 feature grids. Firstly, we generate the molecular surface and then select 800 positive samples (green-colored surface points) from the binding site and 800 negative samples (red-colored surface points) randomly from non-binding regions of the surface, based on the 4.5 °A distance threshold. For each of the 1,600 sampled points, we construct a 41×41×41×22 feature grid, which includes geometric, chemical and force-field information. Grids corresponding to binding site surface points are labeled as 1 (green cube), while grids corresponding to non-binding surface points are labeled as 0 (red cube).

**Fig. 1.**
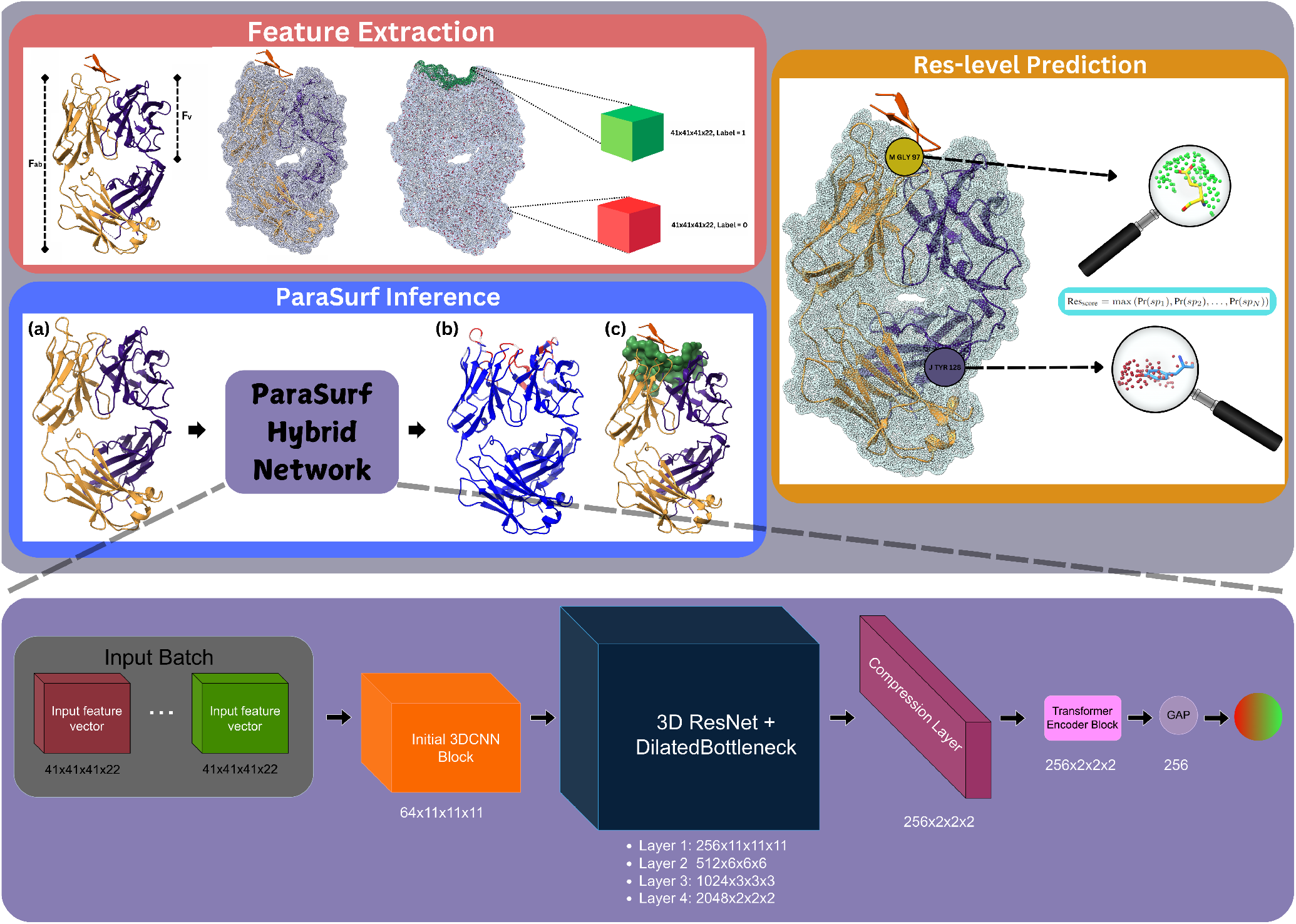
Overview of the ParaSurf framework for paratope prediction. The top section (Feature Extraction) illustrates the Feature Extraction process, capturing molecular surface points for geometric, chemical and force-field data to build grid-based inputs. The middle section (ParaSurf Inference) represents the ParaSurf Inference stage, where the ParaSurf hybrid deep learning network processes the antibody as input (a) to predict binding regions (b, c). The right side (Res-level Prediction) demonstrates Residue-Level Prediction, showing the predicted binding sites of two selected residues (GLY and TYR). The bottom section outlines the ParaSurf network architecture.

In the Res-level Prediction section of Figure 1 we illustrate how residues are classified as binding or non-binding. Using the same PDB complex “5EOC,” we focus on two residues. The first is a Glycine (GLY) residue from the light chain “M” and is the 97th residue in that chain. As shown in the image, GLY belongs to the binding site. The second is a Tyrosine (TYR) residue from the heavy chain “J,” specifically the 128th residue of that chain, which does not belong to the binding site. To determine the score for each residue, we apply Equation 1. The residue score for GLY is calculated as the maximum prediction score from the surrounding surface points of that particular residue (highlighted in green). Similarly, the residue score for TYR is determined by the maximum prediction score of its surrounding surface points (highlighted in red). If the residue score exceeds a certain threshold, the residue is classified as binding.

In the ParaSurf Inference section of Figure 1, we observe how a “blind” inference is performed. ParaSurf takes as input only the paratope (without any antigen information), shown in 1(a) and outputs the predicted binding sites (shown in red). The intensity of the red color corresponds to the prediction confidence, with darker shades indicating a higher likelihood of the residue belonging to the binding site. We follow the common convention of annotating the predicted probabilities for each residue in the b-factor column of the original paratope PDB structure (last column of the PDB file), depicted in 1(b). Additionally, we generate a “pocket” PDB file that highlights the predicted surface points of the binding site (green surface in 1(c)).

Prior works, including DeepSurf2.0 (Papadopoulos et al. 2023) and DeepSurf2-FF (Papadopoulos et al. 2024), laid the foundation for ParaSurf. These studies focused on surface-level binding site prediction, emphasizing on paratope surface points utilizing simpler 3D ResNet architectures. While DeepSurf2.0 demonstrated the effectiveness of geometric and chemical feature integration, DeepSurf2-FF advanced this by introducing electrostatic force fields, further enhancing the predictive power for antibody-binding site predictions. These preliminary results, showcased the model’s potential for surface-level predictions and provided the motivation to develop the more sophisticated ParaSurf framework.

##### Algorithm 1

ParaSurf Feature Extraction

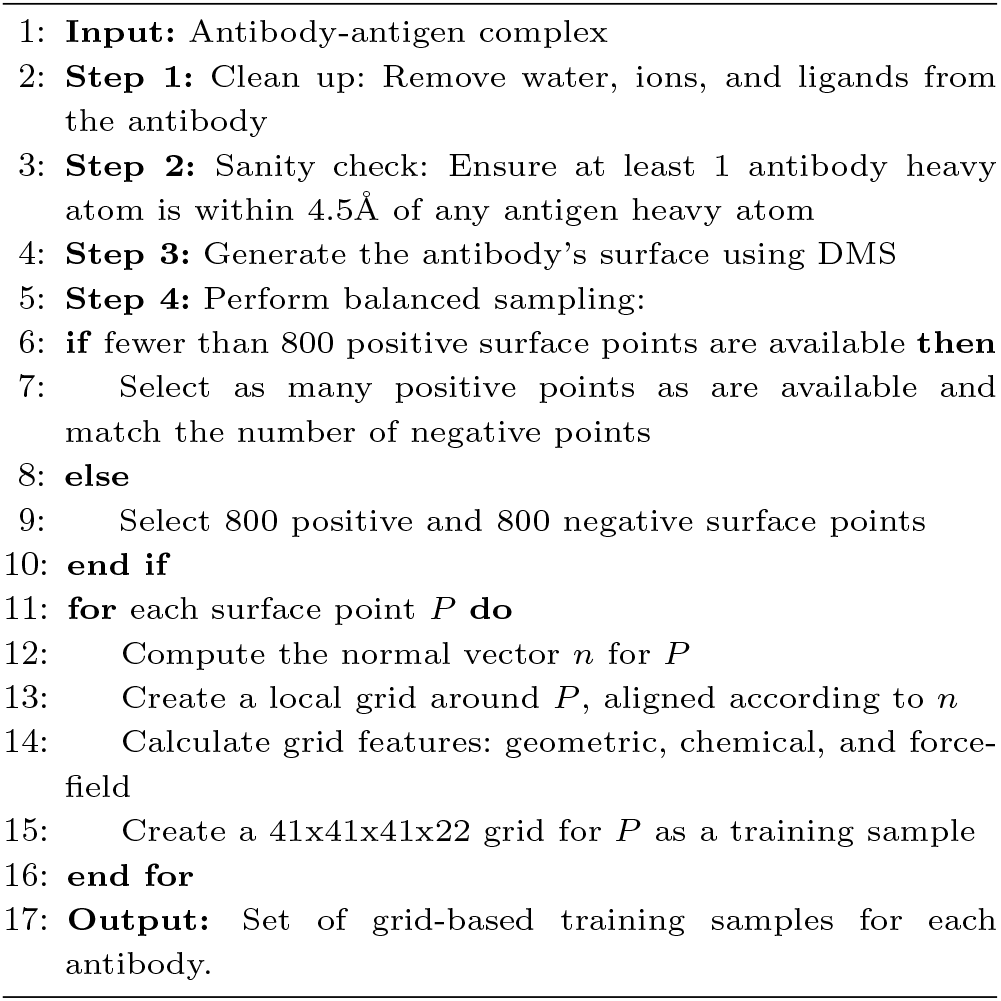

Before conducting the main experiments on the antibody-antigen datasets, we performed a preliminary training round on the DBv5, a general-purpose protein-protein interaction dataset, consisting of 230 complexes. This step was carried out exclusively to fine-tune important parameters for our model, such as the selection of 800 surface points, the grid size of 41×41×41, and the inclusion of electrostatic force field features (AMBER, CHARMM). The goal was to determine the optimal feature extraction and model parameters. No model weights were carried over from this process, as the preliminary training was solely intended for parameter optimization and not for developing the final model. Further details can be found in the Supplementary Information.

### ParaSurf Netwok

ParaSurf employs a hybrid deep learning approach that combines 3D CNN and a transformer block enabling the model to capture local spatial features and global contextual relationships between surface points.

The network begins with an initial 3D CNN block that processes the input 41×41×41×22 tensor, extracting low-level spatial features. Following this, the data is passed through four residual layers composed of dilated bottleneck blocks. These residual layers are inspired by ResNet (He et al. 2016), designed to address challenges like vanishing gradients and performance degradation in deep networks. The inclusion of dilated bottlenecks allows the network to capture multi-scale spatial dependencies. After the residual layers, a compression layer (with a 1×1×1 kernel) reduces feature dimensionality from 2048 channels to 256, lowering the computational cost of the subsequent layers while retaining important information. The output is then batch-normalized and activated using ReLU. Next, the model applies a transformer block, which focuses on learning long-range dependencies between surface points. This block, equipped with self-attention mechanisms (Vaswani 2017) enables the network to focus on relevant parts of the input, capturing how different surface points interact. After the transformer block, global average pooling (GAP)is applied to summarize the learned features, reducing the spatial dimensions and condensing the information into a single vector per 3D grid. A dropout layer (with a dropout rate of 0.1) is applied to prevent overfitting. Finally, the resulting features are passed through a fully connected layer, generating a binding score between 0 and 1 for each surface point, indicating whether a residue belongs to the binding site or not, based on the predicted scores of its surface points. The ParaSurf architecture is illustrated in Figure 1 (bottom). The input to the model consists of batches of 41×41×41×22 grids, where green represents binding grids and red represents non-binding grids. The model processes these inputs and outputs a binding score between 0 and 1, as indicated by the gradient ball, transitioning from red (non-binding) to green (binding).

### Training and Evaluation

#### Training

ParaSurf framework was implemented in Pytorch, in an NVIDIA RTX 3090 GPU. The network was trained using binary cross-entropy loss with the Adam optimizer. A learning rate scheduler was applied, starting at 1e−4 and decaying by a factor of 5 every 5 epochs to facilitate gradual convergence. The batch size was set to 64 for all training scenarios. Full training details, including parameter tuning and adjustments, are provided in the Supplementary Information. Due to incompatibilities with the pdb2pqr software, a few PDB structures were excluded from the training set of each dataset. Additionally, structures that failed the cleaning step, as described earlier, were also removed. The details of these exclusions can be found in the Supplementary Information.

#### Evaluation

ParaSurf’s performance was primarily validated using **AUC-PR** and **AUC-ROC**, which align with the main evaluation metrics reported by previous state-of-the-art methods. **AUC-PR** is particularly useful for assessing performance on imbalanced datasets by focusing on the minority class (binding site residues), offering insights into precision and recall trade-offs, while **AUC-ROC** measures the model’s ability to discriminate between binding and non-binding atoms. Although prior works primarily highlighted these two metrics, they also reported **F-score** and **MCC** (Matthews Correlation Coefficient). F-score is used to balance the importance of precision and recall and indicates the predictive performance, and MCC offers a reliable statistical rate as a measure of the quality of binary classifications. To secure a thorough evaluation, we have included these additional metrics.

To ensure a fair comparison with previous methods, ParaSurf provides results for three evaluation scenarios **ParaSurf(CDR±2), ParaSurf(Fv)** and **ParaSurf(Fab)**. These scenarios allow for both direct comparison with existing state-of-the-art methods and a demonstration of ParaSurf’s ability to generalize across the entire Fab region.

ParaSurf(CDR±2) focuses specifically on the CDR±2 region, directly aligning with methods such as Paragraph and Parapred (Liberis et al. 2018), which concentrate on predicting binding sites within this region. Although Paragraph and Parapred extend their predictions to the entire Fv region by assigning a score of zero to residues outside the CDR±2 loops, ParaSurf provides a direct evaluation of its performance both within the CDR±2 loops and across the full Fv region. As a result, both ParaSurf(CDR±2) and ParaSurf(Fv) serve as direct comparison points for assessing performance against these methods, allowing for a more comprehensive evaluation across both focused and extended regions.

For comparison with methods like Daberdaku (Daberdaku and Ferrari 2019, PECAN (Pittala and Bailey-Kellogg 2020) and MIPE (Wang et al. 2024), which predict binding sites across the entire Fv region, we present results as ParaSurf(Fv). Unlike methods that extend predictions from the CDR±2 region to the full Fv region using zero-padding, ParaSurf makes predictions across the Fv region directly, without the need for extrapolation, offering a more direct assessment of binding site prediction over the entire Fv.

As such, ParaSurf(Fv) serves as a comprehensive comparison point against all methods, encompassing both specific CDR±2 predictions and full Fv region evaluations.

As a significant contribution of this work, we also introduce ParaSurf(Fab), which extends binding site predictions across the entire Fab region of the antibody. This evaluation not only surpasses the Fv region but also highlights ParaSurf’s capacity to predict binding sites on a larger scale, demonstrating its ability to generalize beyond traditional regions.

We use the IMGT numbering system (Lefranc et al. 2009) for direct comparison with previous works, where the regions of the variable domain of an antibody are defined as follows: FR1 (residues 1–26), CDR1 (27–38), FR2 (39–55), CDR2 (56–65), FR3 (66–104), CDR3 (105–117), and FR4 (118–128). In this context, FR refers to the framework region, while CDR represents the complementarity-determining region. The Fv region encompasses both FR and CDR regions. To provide a thorough validation of each paratope, ParaSurf was evaluated at three levels: ParaSurf(CDR±2), ParaSurf(Fv), and ParaSurf(Fab) in ascending order.

## Results

In this section, we present the results of ParaSurf across three antibody-antigen interaction prediction benchmarks: PECAN, Paragraph-expanded, and the MIPE dataset. For a comprehensive evaluation, we report results for ParaSurf at three levels: ParaSurf(CDR±2), ParaSurf(Fv), and ParaSurf(Fab). Each evaluation is conducted using the standard metrics: AUC-PR, AUC-ROC, F1 score, and MCC. Furthermore, in alignment with the thresholds used in the Paragraph model, we evaluate ParaSurf at 0.5 and 0.734 thresholds, the latter being used for direct comparison with the Paragraph model, while the 0.5 threshold remains the most commonly used across similar tasks.

In addition, we also included a baseline method similar to that used by Paragraph (Chinery et al. 2023). This baseline measures the proportion of residues that bind the antigen at each sequence position in the training set and uses these proportions to predict binding in the test set. As the original MIPE paper did not provide a baseline, we calculated this for the MIPE dataset, ensuring that all benchmarks are evaluated using the same reference point. This provides a useful benchmark to understand the improvements achieved by model-driven predictions.

### PECAN Dataset

The PECAN dataset is widely recognized as a benchmark for evaluating paratope prediction models, consisting of 460 antibody-antigen complexes. Table 1 shows the comparison of various models, including the baseline and previous state-of-the-art methods, along with the performance of ParaSurf across different levels and thresholds.

**Table 1.** Comparison of Paratope Prediction Methods on the PECAN Dataset. ParaSurf is evaluated at both 0.5 and 0.734 thresholds. The best results for each metric at the 0.734 threshold are highlighted.

| Method | PR AUC | ROC AUC | MCC | F1 |
| --- | --- | --- | --- | --- |
| Baseline | 0.626 | 0.952 | 0.635 | 0.665 |
| Daberdaku | 0.545 | 0.923 | - | - |
| Parapred | 0.646 | 0.930 | - | - |
| PECAN | 0.675 | 0.952 | - | - |
| Paragraph | 0.696 | 0.934 | **0.654** | **0.685** |
| ParaSurf (CDR±2) | 0.739 / 0.739 | 0.952 / 0.952 | 0.561 / 0.592 | 0.600 / 0.638 |
| ParaSurf (Fv) | 0.733 / **0.733** | 0.955 / **0.955** | 0.584 / 0.612 | 0.608 / 0.647 |
| ParaSurf (Fab) | 0.730 / 0.730 | 0.961 / 0.961 | 0.595 / 0.614 | 0.611 / 0.648 |

We observe a clear improvement in performance as ParaSurf moves from the CDR±2 region to the full Fab region. This trend is evident across all evaluation metrics—PR AUC, ROC AUC, MCC, and F1 score. The consistent increase underscores the robustness of ParaSurf and its ability to generalize predictions across the entire antibody structure. This not only highlights the model’s capability to capture binding residues effectively within the core CDR±2 region but also its potential to predict binding interactions over a wider region of the antibody, including the framework (FR) regions.

By directly comparing ParaSurf(Fv) with Paragraph (highlighted in the table), we see that ParaSurf achieves consistent improvements in the key metrics of AUC-ROC and AUC-PR, further confirming the effectiveness and generalizability of the model across the antibody regions. While ParaSurf shows marginal differences in MCC and F1, its primary strength lies in robust discrimination, as captured by the AUC metrics, which remain the core focus for evaluating model performance.

### Paragraph Expanded Dataset

Table 2 provides a comparison between ParaSurf and the Paragraph model on the Paragraph-expanded dataset, which is a dataset consisting of 1,086 antibody-antigen complexes. Similar to PECAN results, ParaSurf shows consistent improvements across all metrics. Notably, ParaSurf(Fv) outperforms Paragraph, particularly in PR AUC, ROC AUC, and F1 scores, confirming its robustness and ability to generalize across the Fv and the Fab regions.

**Table 2.** Comparison of Paratope Prediction Methods on the Paragraph Expanded Dataset. ParaSurf is evaluated at both 0.5 and 0.734 thresholds. The best results for each metric at the 0.734 threshold are highlighted.

| Method | PR AUC | ROC AUC | MCC | F1 |
| --- | --- | --- | --- | --- |
| Baseline | 0.624 | 0.952 | 0.654 | 0.622 |
| Paragraph | 0.725 | 0.934 | **0.696** | 0.669 |
| ParaSurf (CDR±2) | 0.787 / 0.787 | 0.960 / 0.960 | 0.604 / 0.655 | 0.626 / 0.683 |
| ParaSurf (Fv) | 0.793 / **0.793** | 0.967 / **0.967** | 0.630 / 0.676 | 0.645 / **0.698** |
| ParaSurf (Fab) | 0.795 / 0.795 | 0.973 / 0.973 | 0.643 / 0.674 | 0.651 / 0.686 |

We also report additional results for the Paragraph expanded dataset in the Supplementary Information for training scenarios 3 and 4, where we trained ParaSurf using solely one chain (either heavy or light) in each case. In these experiments, ParaSurf consistently achieved state-of-the-art results compared to Paragraph. Furthermore, we conducted a detailed CDR loop analysis, reporting performance across the four key metrics for each CDR loop and the Framework region.

Notably, ParaSurf demonstrates exceptional performance on the most variable CDRH3 loop, achieving an AUC-ROC of 0.959 and an AUC-PR of 0.895, compared to Paragraph’s 0.866 and 0.796, respectively. Given the high variability of CDRH3, this level of precision is crucial for accurately identifying paratope binding sites in the most dynamic region of the antibody, where variability often leads to diverse antigen recognition patterns. Similar improvements are observed in the CDRL3 variable loop, where ParaSurf achieved AUC-ROC 0.989 and an AUC-PR of 0.910, compared to Paragraph’s 0.884 and 0.770, respectively Additionally, ParaSurf excels at predicting binding residues in the Framework region, where class imbalance is a known challenge due to the small proportion of framework residues contributing to paratope binding. With our balanced sampling technique during feature extraction, ParaSurf mitigates the inherent class imbalance issue and generalizes effectively across the Framework region, achieving an AUC-ROC of 0.981 and an AUC-PR of 0.805. In contrast, Paragraph struggles with this imbalance, reporting only 0.768 and 0.429, respectively. For all this analysis refer to Supplementary Information.

### MIPE Dataset

The MIPE dataset is a relatively newer benchmark for paratope prediction, consisting of 626 antibody-antigen complexes. Table 3 presents a comparison of the baseline and various models, including the performance of ParaSurf across different levels and thresholds. Since the original MIPE paper did not specify a clear classification threshold, we report results for both the 0.5 and 0.734 thresholds to ensure a consistent comparison across methods. Additionally, we calculated a baseline for the MIPE dataset, similar to the one used for the PECAN and Paragraph datasets, to provide a common reference for evaluation.

**Table 3.** Comparison of Paratope Prediction Methods on the MIPE Dataset. ParaSurf is evaluated at both 0.5 and 0.734 thresholds. The best results for each metric at the 0.734 threshold are highlighted.

| Method | PR AUC | ROC AUC | MCC | F1 |
| --- | --- | --- | --- | --- |
| Baseline | 0.465 | 0.931 | 0.177 | 0.536 |
| Parapred | 0.652 | 0.868 | 0.503 | - |
| AG-Fast-Parapred | 0.612 | 0.883 | 0.548 | - |
| PECAN | 0.713 | 0.915 | 0.558 | - |
| Paragraph | 0.650 | 0.927 | 0.488 | 0.616 |
| Pesto | 0.721 | 0.856 | 0.433 | 0.611 |
| MIPE | 0.741 | 0.927 | 0.554 | 0.627 |
| MIPE(AlphaFold2) | 0.723 | 0.910 | 0.531 | 0.617 |
| ParaSurf (CDR±2) | 0.778 / 0.778 | 0.962 / 0.962 | 0.557 / 0.648 | 0.581 / 0.686 |
| ParaSurf (Fv) | 0.781 / **0.781** | 0.967 / **0.967** | 0.581 / **0.659** | 0.597 / **0.690** |
| ParaSurf (Fab) | 0.782 / 0.782 | 0.972 / 0.972 | 0.593 / 0.644 | 0.618 / 0.666 |

ParaSurf outperforms all previous methods across every region of the antibody and at both thresholds, demonstrating its robustness and generalizability also on MIPE dataset.

We report results for all key metrics across all training scenarios for each of the three datasets at the Supplementary Information. In addition to the metrics already discussed, these include Accuracy, Precision, Recall, CAUROC which is the median of aur-roc values offers resilience to the outliers, Negative Predicted Value (NPV), Specificity (SPC), and False Positive Rate (FPR), providing a comprehensive evaluation of ParaSurf’s performance.

### Unified Dataset Performance

ParaSurf achieves the highest overall performance when trained on the unified ParaSurf dataset, which combines PECAN, Paragraph-expanded, and MIPE datasets, yielding an AUC-ROC of 0.9739 and an AUC-PR of 0.8234 (Supplementary Information).

### Qualitative results

Figure 2 presents the predicted results for the antibody-antigen complex with PDB code “1H0D”, which is a part of the Paragraph expanded test set. On the left, we visualize the actual output of the ParaSurf model when only the antibody is provided as input. The model assigns prediction scores to each residue, reflected as b-factor replacements, color-coded by their binding potential. In the middle, we observe confusion matrices summarizing the prediction outcomes at three levels: the entire Fab region, which contains the total amount of the antibody’s residues, the Fv region, and the CDR±2 region. Each matrix shows the number of true negatives (TN), false positives (FP), false negatives (FN), and true positives (TP) residues. On the right, a surface representation highlights the predicted binding site of the paratope, providing a clear view of the specific “pocket” involved in antigen binding.

**Fig. 2.**
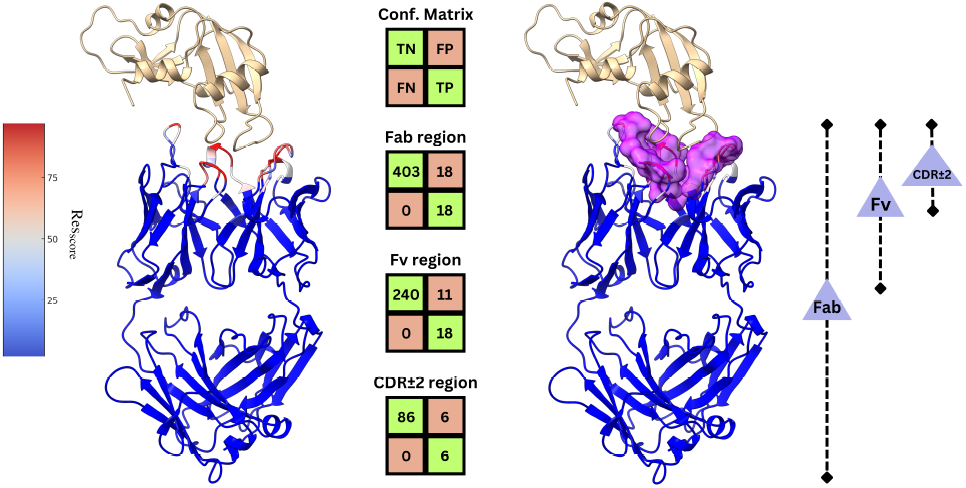
Predicted binding site results for the 1H0D antibody-antigen complex, showing confusion matrices for the Fab, Fv, and CDR±2 regions, alongside a surface visualization of the predicted paratope binding site.

## Discussion

Overall, ParaSurf’s comprehensive evaluation across different thresholds, antibody regions, and diverse datasets underscores its robustness, flexibility, and generalizability. By addressing key challenges such as the inherent class imbalance nature of the paratope prediction task and prediction across larger structural regions like Fab, ParaSurf sets a new standard for paratope prediction. In addition to its general performance, ParaSurf demonstrates exceptional results in highly variable regions of the antibody, particularly the CDRH3 and CDRL3 loops. CDRH3 is the most variable and functionally important loop in antigen binding, as it undergoes significant diversification during somatic recombination—a process critical to generating antibody specificity. The ability to accurately predict binding sites within these regions, especially the CDRH3 loop, enhances ParaSurf’s relevance in antibody design and its capacity to capture key antigen recognition features.

By offering a more complete picture of antibody-antigen interactions, ParaSurf provides a powerful tool for advancing antibody engineering and understanding immune responses at the molecular level. Its outstanding performance in the CDRH3 region, where specificity and antigen recognition are largely determined, highlights the model’s contribution to improving antibody targeting accuracy (Gabrielli et al. 2009).

Furthermore, ParaSurf lays a strong foundation for future work aimed at integrating its predictions with antibody-antigen docking tools. By focusing on the predicted interaction binding surface rather than the entire protein structure, this approach has the potential to significantly enhance docking accuracy while reducing computational time. Ultimately, this integration will improve the efficiency and precision of docking simulations, further supporting advancements in therapeutic antibody development.

## Supporting information

ParaSurf_Supplementary_Information

