## Supplementary material for "ParaSurf: A Surface-Based Deep Learning Approach for Paratope-Antigen Interaction Prediction": ParaSurf_Supplementary_Information

### 1. Datasets

The data-separation was made to ensure that paratopes shared no more than 95% pairwise sequence identity, by using CD-HIT(Li and Godzik (2006)). It includes only complexes with paired heavy and light chains, with a resolution below 3Å and protein antigens. A residue is considered part of the paratope's binding site if any of its heavy atoms (non-hydrogen atoms) is located within **4.5Å** of any antigen-heavy atom.

#### 1.1. PECAN Dataset

The PECAN dataset (Pittala and Bailey-Kellogg (2020)) consists of 460 paratope-antigen complexes, with 205 allocated for training, 103 for validation, and 152 for testing.

ParaSurf begins with a clean-up filtering step to remove ions, small molecules, and ligands from each paratope. This is followed by feature extraction using the OpenBabel module (O'Boyle et al. (2008)) and pdb2pqr (Dolinsky et al. (2004)) to obtain the necessary chemical and electrostatic features respectively. Due to incompatibilities with these modules—such as missing structural data, parsing errors, or other technical issues—some antibody-antigen complexes were excluded from the dataset. Additionally, as noted in the Paragraph supplementary material, three more complexes were removed due to misplacement of the antigen relative to its corresponding antibody. After these adjustments, the final dataset comprised:

- **Training**: 195 complexes,
- **Validation**: 101 complexes, and
- **Test**: 152 complexes.

In total, 12 complexes were removed from the original PECAN dataset.

#### 1.2. Paragraph Expanded Dataset

The Paragraph Expanded Dataset (Chinery et al. (2023)) consists of 1,086 antibody-antigen complexes, which are divided into training, validation, and test sets using a 60-20-20 split, following the exact same data partitioning as in the original Paragraph study. This results in 651 complexes for training, 217 for validation, and 218 for testing. After applying the filtering and feature extraction procedures, the final dataset included:

- **Training**: 626 complexes,
- **Validation**: 214 complexes, and

© .

- **Test**: 216 complexes.

In total, 30 complexes were removed from the original Paragraph Expanded Dataset.

#### 1.3. MIPE Dataset

The MIPE Dataset (Wang et al. (2024)) consists of 626 antibody-antigen complexes. Following the same methodology as the original MIPE study, we randomly split the dataset, allocating 90% (563 complexes) for cross-validation (CV) and 10% (63 complexes) as the same independent test set. After applying the filtering and feature extraction procedures, the final dataset comprised:

- **Training-Validation**: 528 complexes for 5-fold CV, and
- **Test**: 63 complexes.

In total, 35 complexes were removed from the original MIPE Dataset.

#### 1.4. ParaSurf Dataset

The ParaSurf dataset is a unified dataset combining three benchmarks: PECAN, Paragraph-expanded, and MIPE. Following a thorough process of filtering and removing overlapping occupancies, the final dataset was split as follows:

- **Training Set:** 1,112 antibody-antigen complexes,
- **Validation Set:** 298 complexes, and
- **Test Set:** 394 complexes.

This dataset integrates diverse antibody-antigen structures from all three benchmarks, providing a robust and comprehensive resource for training and evaluating models like ParaSurf.

### 2. Finetune ParaSurf Parameters

To optimize ParaSurf's performance during the feature extraction and training phases, we conducted a preliminary training round using the Docking Benchmark Version 5 (DBv5) dataset (Vreven et al. (2015)), which is a general-purpose protein-protein interaction dataset consisting of 230 complexes.

It is crucial to note that no model weights were retained from this process, as the objective was solely parameter tuning.

The parameters finetuned included grid input size, chemical and force-field features, number of training samples, and various training hyperparameters. We also experimented with different model architectures and pooling methods. The specific parameters that yielded the best results were as follows:

**Feature Extraction:**

- **Grid size**: 41x41x41 grid
- **Chemical features**: 18 chemical descriptors

- **Force-field features**: 2 force-field descriptors (AMBER and CHARMM) and 2 atom radii features
- **Number of samples**: ±800 positive and negative samples per complex

**Training Parameters:**

- **Learning rate (lr)**: An lr scheduler was applied, starting at 1e4 and decaying by a factor of 5 every 5 epochs
- **Batch size**: 64
- **Dropout**: 10%
- **Model architecture**: 3D ResNet combined with a transformer layer
- **Pooling method**: Global Average Pooling (GAP).

### 3. ParaSurf evaluation

In this section, we present a comprehensive evaluation of ParaSurf across three antibody-antigen interaction benchmarks. For each dataset, we provide detailed performance metrics across different training scenarios, complemented by visualizations that illustrate the model's predictive capabilities. Each training scenario is discussed in its own subsection, offering a thorough analysis of ParaSurf's behavior under varying conditions.

The key validation metrics used for this evaluation are AUC-PR and AUC-ROC, which are the most critical indicators of performance, particularly for imbalanced datasets. Additionally, we report MCC, F1, Accuracy (Acc), Precision (Pr), Recall (Rc), and CAUROC—a median of AUC-ROC values that is resilient to outliers. To further ensure a comprehensive evaluation, we include Negative Predicted Value (NPV), Specificity (SPC), and False Positive Rate (FPR).

For each scenario, we report results for ParaSurf at three levels: ParaSurf(CDR±2), ParaSurf(Fv), and ParaSurf(Fab) using two thresholds—0.5 and 0.734 respectively (with the latter specifically recommended for direct comparison with the Paragraph model).

#### 3.1. Scenario 1

In this scenario ParaSurf is trained and tested on the most prominent PECAN dataset. Results are shown below in table 1.

In the image S1, we provide two examples from the PECAN test set to illustrate the ParaSurf inference process. On the left side of each example, ParaSurf outputs the predicted binding sites (shown in red). The intensity of the red color corresponds to the prediction confidence, with darker shades indicating a higher likelihood of the residue belonging to the binding site. We follow the common convention of annotating the predicted probabilities for each residue in the b-factor column of the original paratope PDB structure (last column of the PDB file). On the right side, we show a ”pocket” PDB file that highlights the predicted surface points of the binding site, giving a clear view of the binding interface.

| **Method** | **PR AUC** | **ROC AUC** | **MCC** | **F1** | **Acc** | **Pr** | **Rc** | **CAUROC** | **NPV** | **SPC** | **FPR** |
|---|---|---|---|---|---|---|---|---|---|---|---|
| Baseline | 0.626 | 0.952 | 0.635 | 0.665 | - | - | - | - | - | - | - |
| Daberdaku | 0.545 | 0.923 | - | - | - | - | - | - | - | - | - |
| Parapred | 0.646 | 0.930 | - | - | - | - | - | - | - | - | - |
| PECAN | 0.675 | 0.952 | - | - | - | - | - | - | - | - | - |
| Paragraph | 0.696 | 0.934 | **0.654** | **0.685** | - | - | - | - | - | - | - |
| ParaSurf (CDR±2) | 0.739 / 0.739 | 0.952 / 0.952 | 0.561 / 0.592 | 0.600 / 0.638 | 0.867 / 0.897 | 0.453 / 0.528 | 0.871 / 0.806 | 0.959 / 0.959 | 0.983 / 0.975 | 0.865 / 0.907 | 0.135 / 0.093 |
| ParaSurf (Fv) | 0.733 / **0.733** | 0.955 / **0.955** | 0.584 / 0.612 | 0.608 / 0.647 | 0.890 / 0.913 | 0.495 / 0.534 | 0.880 / 0.821 | 0.969 / 0.969 | 0.987 / 0.980 | 0.889 / 0.922 | 0.111 / 0.078 |
| ParaSurf (Fab) | 0.730 / 0.730 | 0.961 / 0.961 | 0.595 / 0.614 | 0.611 / 0.648 | 0.907 / 0.923 | 0.467 / 0.513 | 0.883 / 0.843 | 0.975 / 0.975 | 0.989 / 0.985 | 0.908 / 0.930 | 0.092 / 0.070 |

Table 1: Comparison of Paratope Prediction Methods on the PECAN test set.

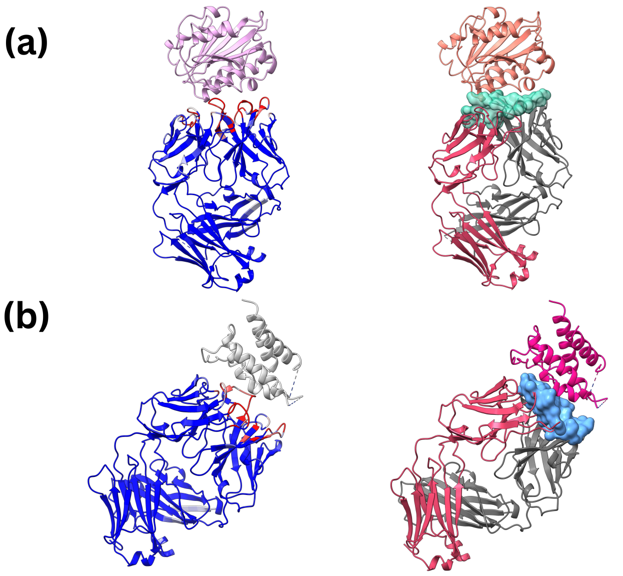


Figure S1: Visualization of ParaSurf results on PECAN test set: (a) shows the predicted binding sites for the 3EOA PDB structure, and (b) shows the predicted binding sites for the 4YUE PDB structure. The left side of each image represents the predicted binding sites, while the right side highlights the binding surface as predicted by ParaSurf.

### 3.2. Scenario 2

In this scenario, ParaSurf is trained and tested on the Paragraph Expanded dataset. Results are shown below in table 2.

In the image S2, we provide two examples from the Expanded Paragraph test set to illustrate the ParaSurf inference process.

Additionally, we report results for each CDR loop and the framework region, directly comparing them with the state-of-the-art Paragraph model, on table 3. For simplicity, we present only the most important metrics (AUC-PR, AUC-ROC, MCC, and F1), using the same threshold of 0.734 as used in the Paragraph evaluation.

| Method | PR AUC | ROC AUC | MCC | F1 | Acc | Pr | Rc | CAUROC | NPV | SPC | FPR |
|---|---|---|---|---|---|---|---|---|---|---|---|
| Baseline | 0.624 | 0.952 | 0.654 | 0.622 | - | - | - | - | - | - | - |
| Paragraph | 0.725 | 0.934 | **0.696** | 0.669 | - | - | - | - | - | - | - |
| ParaSurf (CDR±2) | 0.787 / 0.787 | 0.960 / 0.960 | 0.604 / 0.655 | 0.626 / 0.683 | 0.884 / 0.920 | 0.499 / 0.626 | 0.887 / 0.797 | 0.972 / 0.972 | 0.983 / 0.970 | 0.884 / 0.937 | 0.116 / 0.063 |
| ParaSurf (Fv) | 0.793 / **0.793** | 0.967 / **0.967** | 0.630 / 0.676 | 0.645 / **0.698** | 0.905 / 0.933 | 0.519 / 0.639 | 0.892 / 0.808 | 0.979 / 0.979 | 0.986 / 0.976 | 0.906 / 0.948 | 0.093 / 0.052 |
| ParaSurf (Fab) | 0.795 / 0.795 | 0.973 / 0.973 | 0.643 / 0.674 | 0.651 / 0.686 | 0.921 / 0.940 | 0.525 / 0.605 | 0.893 / 0.837 | 0.984 / 0.984 | 0.989 / 0.982 | 0.923 / 0.951 | 0.077 / 0.049 |

Table 2: Comparison of Paratope Prediction Methods on the Paragraph Expanded test set.

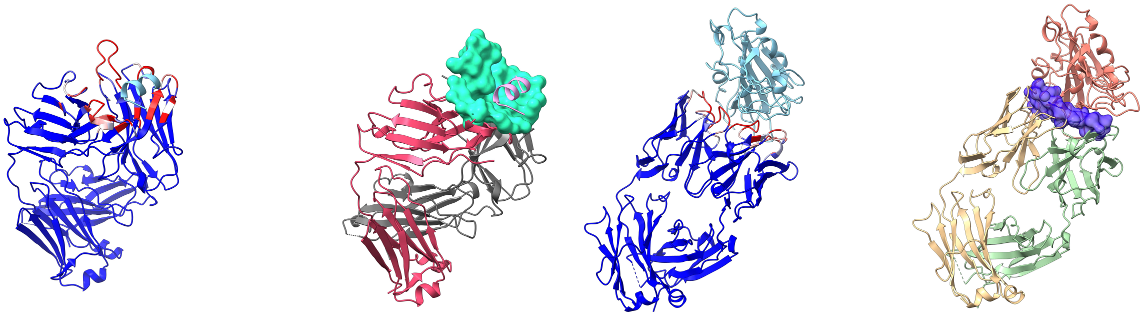

**(a)** **(b)**

Figure S2: Visualization of ParaSurf results on Paragraph Expanded test set: (a) shows the predicted binding sites for the 4N0Y PDB structure, and (b) shows the predicted binding sites for the 7S0B PDB structure. The left side of each image represents the predicted binding sites, while the right side highlights the binding surface as predicted by ParaSurf.

Notably, ParaSurf demonstrates outstanding generalization across the entire Fab domain, particularly in the framework region. Unlike Paragraph, which struggles to generalize beyond the CDR±2 region, ParaSurf significantly improves prediction performance in the framework region, substantially outperforming both the baseline and Paragraph models. This improvement indicates ParaSurf's ability to manage the extreme class imbalance typically found in the framework region, where paratope residues are sparse.

Equally important is ParaSurf's superior performance in the most variable regions of the antibody. In the CDR-H3 loop, which is crucial for antigen binding due to its high variability, ParaSurf achieved state-of-the-art results. Its ability to excel in this region, as well as the CDR-L3 loop, further highlights ParaSurf's robustness in modeling complex regions that play a critical role in antigen-antibody interactions. Results for the CDR loop analysis are shown below in table 3.

| Method | CDR-L1 | | | |
|---|---|---|---|---|
| | PR AUC | ROC AUC | F-score | MCC |
| Baseline | 0.691 | 0.799 | 0.642 | 0.435 |
| Paragraph | 0.762 | 0.857 | 0.678 | 0.499 |
| ParaSurf | **0.885** | **0.965** | **0.692** | **0.669** |
| **Method** | **CDR-L2** | | | |
| | PR AUC | ROC AUC | F-score | MCC |
| Baseline | 0.238 | 0.741 | 0.555 | 0.429 |
| Paragraph | 0.675 | 0.876 | **0.654** | **0.559** |
| ParaSurf | **0.772** | **0.932** | 0.535 | 0.509 |
| **Method** | **CDR-L3** | | | |
| | PR AUC | ROC AUC | F-score | MCC |
| Baseline | 0.682 | 0.827 | 0.716 | 0.545 |
| Paragraph | 0.770 | 0.884 | **0.747** | 0.598 |
| ParaSurf | **0.910** | **0.989** | 0.617 | **0.619** |
| **Method** | **CDR-H1** | | | |
| | PR AUC | ROC AUC | F-score | MCC |
| Baseline | 0.514 | 0.810 | 0.650 | 0.468 |
| Paragraph | 0.735 | 0.856 | 0.678 | 0.515 |
| ParaSurf | **0.868** | **0.964** | **0.681** | **0.652** |
| **Method** | **CDR-H2** | | | |
| | PR AUC | ROC AUC | F-score | MCC |
| Baseline | 0.643 | 0.745 | 0.667 | 0.368 |
| Paragraph | 0.789 | 0.854 | 0.727 | 0.498 |
| ParaSurf | **0.896** | **0.963** | **0.748** | **0.710** |
| **Method** | **CDR-H3** | | | |
| | PR AUC | ROC AUC | F-score | MCC |
| Baseline | 0.700 | 0.813 | 0.735 | 0.519 |
| Paragraph | 0.796 | 0.866 | **0.762** | 0.571 |
| ParaSurf | **0.895** | **0.959** | 0.759 | **0.709** |
| **Method** | **Framework** | | | |
| | PR AUC | ROC AUC | F-score | MCC |
| Baseline | 0.410 | 0.952 | 0.436 | 0.427 |
| Paragraph | 0.429 | 0.768 | 0.505 | 0.500 |
| ParaSurf | **0.805** | **0.981** | **0.723** | **0.717** |

Table 3: Comparison of PR AUC, ROC AUC, F-score, and MCC across CDR and the Framework regions on the Expanded Paragraph Dataset.

#### 3.3. Scenario 3 and 4

In this scenario, ParaSurf is trained and tested on the Paragraph Expanded dataset, focusing on either the heavy or light chain of the antibody while discarding the other. To ensure valid antigen pairs (fulfilling the 4.5Å Euclidean distance condition), some complexes were removed, resulting in adjusted datasets for these two scenarios.

Training ParaSurf on individual chains (either heavy or light) is crucial for addressing real-world cases where data for only one chain is available. This allows ParaSurf to generalize its predictions even when the paired information from both chains is missing. Previous state-of-the-art methods, such as Paragraph, have demonstrated that performance drops when predicting paratope residues from only one chain, as both chains typically provide complementary information during antigen binding. However, training on single-chain data is essential for models to adapt to situations where only partial information is present.

It is also important to note that the light chain generally faces a greater class imbalance compared to the heavy chain, with fewer residues contributing to the paratope binding site. This imbalance negatively affects performance, as shown in previous works like Paragraph and Parapred. Therefore, these two scenarios test ParaSurf's ability to handle such class imbalance effectively. As expected, neither individual chain's performance is expected to exceed that of paired data, but by training on only heavy or light chains, ParaSurf aims to achieve a competitive performance while mitigating the challenges posed by class imbalance.

#### 3.3.1. SCENARIO 3

In this scenario, ParaSurf is trained and tested on the Paragraph Expanded dataset, using only the heavy chain of the antibody-antigen complexes. Notably, no complexes were removed during this process, as there was always at least one heavy atom from the heavy chain within a 4.5Å Euclidean distance from the antigen atoms. This reinforces the critical role of the heavy chain in antigen binding, highlighting its dominant contribution to the interaction process.

The results of ParaSurf, trained and tested using only the heavy chain, are shown in Table 4. Despite being trained on just one chain, ParaSurf demonstrates robust predictive performance, achieving high AUC-ROC and AUC-PR values across all evaluation levels (CDR±2, Fv, Fab).

| Method | PR AUC | ROC AUC | MCC | F1 | Acc | Pr | Rc | CAUROC | NPV | SPC | FPR |
|---|---|---|---|---|---|---|---|---|---|---|---|
| Baseline | 0.640 | 0.951 | 0.679 | 0.642 | - | - | - | - | - | - | - |
| Paragraph | 0.737 | 0.943 | **0.713** | 0.681 | - | - | - | - | - | - | - |
| ParaSurf (CDR±2) | 0.774 / 0.774 | 0.945 / 0.945 | 0.600 / 0.629 | 0.639 / 0.674 | 0.847 / 0.881 | 0.518 / 0.595 | 0.913 / 0.839 | 0.957 / 0.957 | 0.977 / 0.962 | 0.834 / 0.890 | 0.166 / 0.110 |
| ParaSurf (Fv) | 0.766 / **0.766** | 0.956 / **0.956** | 0.615 / 0.647 | 0.639 / **0.697** | 0.873 / 0.902 | 0.512 / 0.591 | 0.916 / 0.850 | 0.970 / 0.970 | 0.983 / 0.972 | 0.866 / 0.911 | 0.134 / 0.089 |
| ParaSurf (Fab) | 0.764 / 0.764 | 0.964 / 0.964 | 0.625 / 0.647 | 0.655 / 0.684 | 0.895 / 0.914 | 0.510 / 0.563 | 0.917 / 0.873 | 0.977 / 0.977 | 0.987 / 0.979 | 0.891 / 0.920 | 0.110 / 0.080 |

Table 4: Comparison of Paratope Prediction Methods on the Paragraph Expanded test set when trained and tested only on the heavy chain.

This scenario highlights the resilience of ParaSurf when data is limited to just one chain, underscoring its potential applicability in real-world situations where complete antibody information is unavailable. The high recall (Rc) values across all levels also indicate ParaSurf's strong ability to capture true positive binding residues even in these single-chain conditions.

In the image S3, we provide two examples from the Expanded Paragraph test set, where each antibody comprises only the Heavy chain, to illustrate the ParaSurf inference process.

#### 3.3.2. SCENARIO 4

In contrast to the heavy chain, 22 complexes were removed in this scenario, as they no longer satisfied the 4.5Å Euclidean distance criterion. Specifically, 14 complexes were removed from the training set, 6 from the validation set, and 2 from the test set. This reduction is indicative of the lesser contribution of the light chain to antigen binding, compared to the heavy chain.

The results of ParaSurf when trained and tested using only the light chain are presented in Table 5. Despite the inherent challenges posed by the light chain—such as a greater class imbalance and

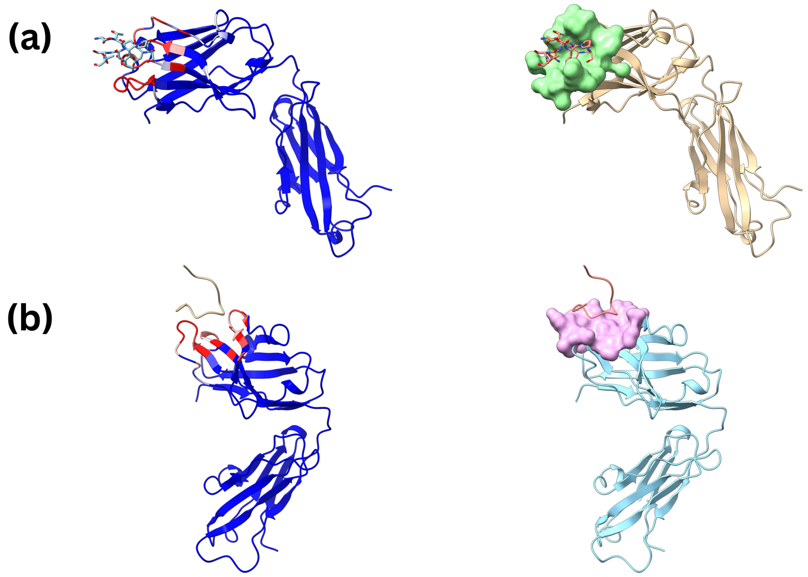


Figure S3: Visualization of ParaSurf results on Paragraph Expanded test set where each antibody comprises of only the Heavy chain: (a) shows the predicted binding sites for the 6VJT PDB structure, and (b) shows the predicted binding sites for the 1FTP PDB structure. The left side of each image represents the predicted binding sites, while the right side highlights the binding surface as predicted by ParaSurf.

| **Method** | **PR AUC** | **ROC AUC** | **MCC** | **F1** | **Acc** | **Pr** | **Rc** | **CAUROC** | **NPV** | **SPC** | **FPR** |
|---|---|---|---|---|---|---|---|---|---|---|---|
| Baseline | 0.603 | 0.949 | 0.612 | 0.584 | - | - | - | - | - | - | - |
| Paragraph | 0.637 | 0.910 | **0.630** | **0.608** | - | - | - | - | - | - | - |
| ParaSurf (CDR±2) | 0.747 / 0.747 | 0.957 / 0.957 | 0.459 / 0.517 | 0.464 / 0.543 | 0.843 / 0.893 | 0.311 / 0.399 | 0.912 / 0.850 | 0.979 / 0.979 | 0.990 / 0.985 | 0.839 / 0.899 | 0.161 / 0.101 |
| ParaSurf (Fv) | 0.728 / **0.728** | 0.963 / **0.963** | 0.488 / 0.541 | 0.488 / 0.560 | 0.866 / 0.906 | 0.332 / 0.415 | 0.917 / 0.860 | 0.979 / 0.979 | 0.992 / 0.987 | 0.863 / 0.912 | 0.136 / 0.088 |
| ParaSurf (Fab) | 0.722 / 0.722 | 0.968 / 0.968 | 0.503 / 0.538 | 0.495 / 0.544 | 0.888 / 0.915 | 0.339 / 0.394 | 0.919 / 0.881 | 0.982 / 0.982 | 0.993 / 0.990 | 0.887 / 0.918 | 0.113 / 0.082 |

Table 5: Comparison of Paratope Prediction Methods on the Paragraph Expanded Dataset test set when trained and tested only on the light chain.

the overall reduced number of binding residues—ParaSurf continues to exhibit robust performance. Particularly notable are the AUC-PR and AUC-ROC, which demonstrate that ParaSurf can still identify binding residues effectively, even in the face of reduced input data. The ability to generalize to the light chain, despite its limitations, indicates the versatility of ParaSurf and its potential to handle diverse antibody-antigen complexes where information is restricted.

In the image S4, we provide two examples from the Expanded Paragraph test set, where each antibody comprises only the light chain, to illustrate the ParaSurf inference process.

#### 3.4. Scenario 5

In this scenario, ParaSurf is trained and tested on the most recent MIPE dataset. Results are shown below in table 6.

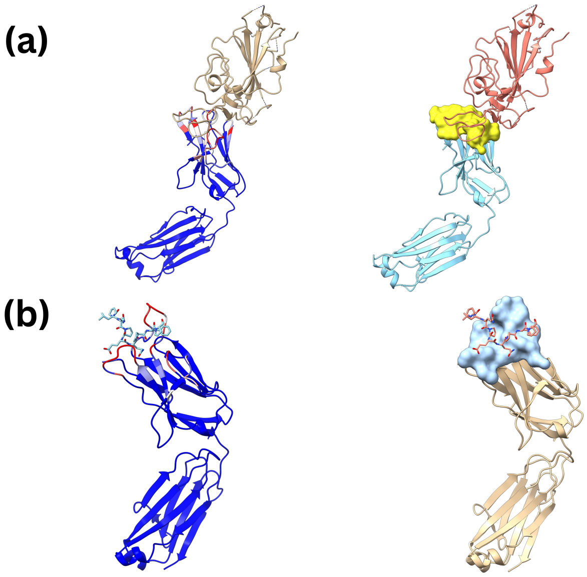


Figure S4: Visualization of ParaSurf results on Paragraph Expanded test set where each antibody comprises of only the Light chain: (a) shows the predicted binding sites for the 7KN4 PDB structure, and (b) shows the predicted binding sites for the 5IJK PDB structure. The left side of each image represents the predicted binding sites, while the right side highlights the binding surface as predicted by ParaSurf.

| **Method** | **PR AUC** | **ROC AUC** | **MCC** | **F1** | **Acc** | **Pr** | **Rc** | **CAUROC** | **NPV** | **SPC** | **FPR** |
|---|---|---|---|---|---|---|---|---|---|---|---|
| Baseline | 0.465 | 0.931 | 0.177 | - | - | - | - | - | - | - | - |
| Parapred | 0.652 | 0.868 | 0.503 | - | - | - | - | - | - | - | - |
| AG-Fast-Parapred | 0.612 | 0.883 | 0.548 | - | - | - | - | - | - | - | - |
| PECAN | 0.713 | 0.915 | 0.558 | - | - | - | - | - | - | - | - |
| Paragraph | 0.650 | 0.927 | 0.488 | - | - | - | - | - | - | - | - |
| Pesto | 0.721 | 0.856 | 0.433 | - | - | - | - | - | - | - | - |
| MIPE | 0.741 | 0.927 | 0.554 | - | - | - | - | - | - | - | - |
| MIPE(AlphaFold2) | 0.723 | 0.910 | 0.531 | - | - | - | - | - | - | - | - |
| ParaSurf (CDR±2) | 0.778 / 0.778 | 0.962 / 0.962 | 0.557 / 0.648 | 0.581 / 0.686 | 0.855 / 0.855 | 0.422 / 0.571 | 0.937 / 0.860 | 0.974 / 0.974 | 0.987 / 0.979 | 0.948 / 0.923 | 0.152 / 0.077 |
| ParaSurf (Fv) | 0.781 / **0.781** | 0.967 / **0.967** | 0.581 / **0.659** | 0.597 / 0.690 | 0.884 / 0.927 | 0.440 / 0.576 | 0.928 / 0.860 | 0.980 / 0.980 | 0.990 / 0.984 | 0.880 / 0.936 | 0.120 / 0.063 |
| ParaSurf (Fab) | 0.782 / 0.782 | 0.972 / 0.972 | 0.593 / 0.644 | 0.617 / 0.667 | 0.909 / 0.933 | 0.464 / 0.536 | 0.918 / 0.881 | 0.984 / 0.984 | 0.991 / 0.988 | 0.908 / 0.939 | 0.092 / 0.061 |

Table 6: Comparison of Paratope Prediction Methods on the MIPE test set.

In the image S5, we provide two examples from the MIPE test set to illustrate the ParaSurf inference process.

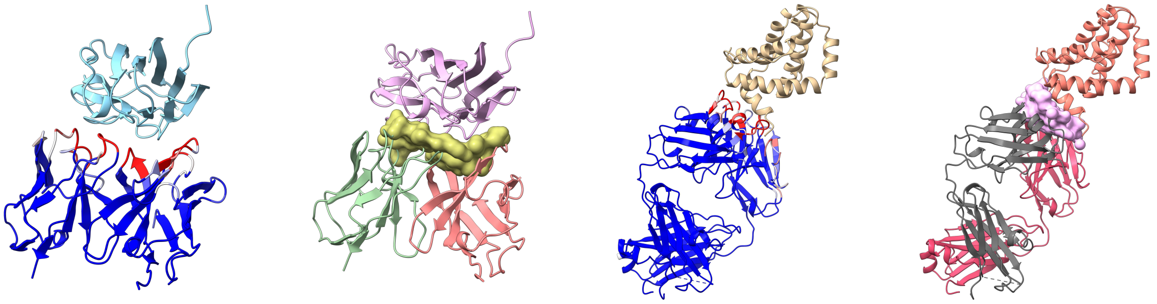

**(a)** **(b)**

Figure S5: Visualization of ParaSurf results on Paragraph Expanded test set: (a) shows the predicted binding sites for the 3AB0 PDB structure, and (b) shows the predicted binding sites for the 5MES PDB structure. The left side of each image represents the predicted binding sites, while the right side highlights the binding surface as predicted by ParaSurf.

### 4. Performance on ParaSurf Dataset

In this scenario, ParaSurf is trained and tested on the ParaSurf Dataset, which is the combination of the dataset (PECAN, Paragraph-expanded, and MIPE). Results are shown below in table 7.

| **Method** | **PR AUC** | **ROC AUC** | **MCC** | **F1** | **Acc** | **Pr** | **Rc** | **CAUROC** | **NPV** | **SPC** | **FPR** |
|---|---|---|---|---|---|---|---|---|---|---|---|
| ParaSurf (CDR±2) | 0.816 / 0.816 | 0.969 / 0.969 | 0.604 / 0.672 | 0.614 / 0.672 | 0.872 / 0.915 | 0.469 / 0.589 | 0.946 / 0.884 | 0.977 / 0.977 | 0.991 / 0.983 | 0.862 / 0.920 | 0.137 / 0.080 |
| ParaSurf (Fv) | 0.823 / **0.823** | 0.974 / **0.974** | 0.630 / 0.693 | 0.634 / **0.707** | 0.895 / 0.930 | 0.490 / 0.606 | 0.945 / 0.888 | 0.982 / 0.982 | 0.993 / 0.986 | 0.890 / 0.935 | 0.110 / 0.065 |
| ParaSurf (Fab) | 0.825 / 0.825 | 0.978 / 0.978 | 0.642 / 0.684 | 0.642 / 0.688 | 0.913 / 0.936 | 0.496 / 0.573 | 0.945 / 0.906 | 0.987 / 0.987 | 0.994 / 0.990 | 0.910 / 0.940 | 0.090 / 0.060 |

Table 7: ParaSurf results on the ParaSurf dataset.

### 5. Overall Results

Figure S6 illustrates the performance of ParaSurf across the three benchmark datasets: PECAN, Paragraph Expanded, and MIPE. The plots compare ParaSurf to other competing methods on the two key metrics: AUC-PR (x-axis) and AUC-ROC (y-axis). Each marker represents a different method, with the combined performance (average of AUC-PR and AUC-ROC) indicated by higher values along both axes. The results show that ParaSurf consistently outperforms competing methods across all benchmarks, achieving higher AUC-PR and AUC-ROC values in each validated region—ParaSurf(CDR±2), ParaSurf(Fv), and ParaSurf(Fab).

### 6. Availability

The source code, the best model weights for each training scenario, the preprocessing pipeline, and the three benchmark datasets with their specific splits are freely available at the Github link: https://github.com/aggelos-michael-papadopoulos/ParaSurf.

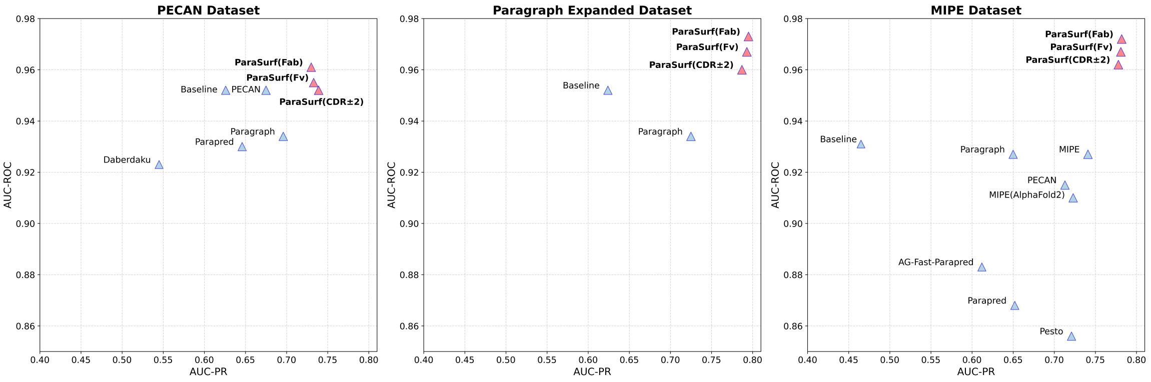


Figure S6: Performance of various methods on the PECAN, Paragraph Expanded, and MIPE datasets. The x-axis represents AUC-PR while the y-axis represents AUC-ROC. Each point corresponds to a method, with the size of the marker indicating combined performance (average of AUC-PR and AUC-ROC). ParaSurf methods are highlighted in bold and marked in red for easy identification.
